# Companion animals harbour globally circulating human-associated *Klebsiella pneumoniae* lineages and high-risk antimicrobial resistance clones

**DOI:** 10.64898/2026.06.13.732079

**Authors:** Stephen Mark Edward Fordham, Elizabeth Sheridan, Francis Drobniewski

## Abstract

Companion animals are increasingly recognised within One Health antimicrobial-resistance (AMR) surveillance, yet the global genomic structure of dog- and cat-associated *Klebsiella pneumoniae* remains poorly defined. We assembled a global dataset of 712 domestic dog- and cat-derived *K. pneumoniae* genomes to define lineage diversity, AMR burden, host-associated patterns and overlap with human-associated populations. Companion-animal isolates comprised 263 sequence types (STs), but recurrent high-risk lineages, including ST307, ST11, ST15 and ST147, were prominent. Among 706 isolates from 25 countries, extended-spectrum β-lactamase (ESBL) and carbapenemase genes were widely distributed, detected in 303 (42.9%) and 98 (13.9%) isolates, respectively. Cat-derived isolates showed higher multidrug-resistance (MDR) prevalence than dog-derived isolates; 80.0% versus 56.3%, partly reflecting enrichment of epidemic clones, especially ST147. MDR was not confined to infection-associated samples, indicating that colonisation may represent an important reservoir state. Comparison with 38,106 human-associated *K. pneumoniae* isolates revealed extensive ST overlap, with 71.1% of companion-animal STs and 87.2% of companion-animal isolates belonging to STs also detected in humans. Focused recombination-filtered phylogenomics of ST147 identified a recent host-spanning MDR sub-lineage linking cat-, dog- and human-associated genomes. Together, these findings show that domestic dogs and cats are not epidemiologically separate from the wider *K. pneumoniae* AMR landscape, but harbour globally circulating human-associated lineages and high-risk AMR clones.

## Introduction

Antimicrobial resistance (AMR) is a major global health challenge that requires coordinated surveillance across human, animal and environmental sectors. The 2025 World Health Organization (WHO) Global Antimicrobial Resistance and Use Surveillance System (GLASS) report highlights the scale of this threat, estimating that one in six laboratory-confirmed bacterial infections caused by common bacterial pathogens in 2023 were resistant to antibiotic treatment [1]. Resistance among Gram-negative pathogens is particularly concerning globally, more than 55% of *Klebsiella pneumoniae* (*K. pneumoniae*) infections were resistant to third-generation cephalosporins (3GCs), a key treatment option for severe infections [1]. These trends reinforce the need for integrated One Health surveillance to identify reservoirs, track transmission pathways and detect clinically important resistant lineages across connected human, animal and environmental settings.

Within this One Health context, companion animals represent an increasingly important but comparatively under-surveilled population. Dogs and cats live in close contact with humans, share household environments and may provide opportunities for bacterial exchange between people, animals and shared surroundings. The scale of this close contact is substantial: in the UK, around 62% of households own a pet, including an estimated 15.5 million dogs and 13 million cats; across Europe there are approximately 90 million dogs and 108 million cats, while the United States has an estimated 87.3 million owned dogs and 76.3 million owned cats [2, 3, 4]. Despite this, companion-animal AMR surveillance remains less developed than human, livestock and food-chain surveillance. In the UK, this gap is now being addressed through Veterinary Medicines Directorate-led initiatives, including a national pilot delivered by Scotland’s Rural College to collect faecal samples from healthy dogs and cats in households, veterinary practices and rescue centres to establish baseline AMR levels and evaluate practical surveillance approaches [5]. This UK system will add to an existing international companion-animal AMR surveillance landscape that includes the China Antimicrobial Resistance Surveillance Network for Pets (CARPet), established in 2021 to monitor clinical bacterial pathogens from companion animals [6].

Companion animals are increasingly recognised as reservoirs of multidrug-resistant (MDR) Enterobacterales, including clinically important *K. pneumoniae* lineages that overlap with human-associated populations. However, the available evidence remains largely based on localised studies, household investigations, outbreak reports, or isolate collections enriched for extended-spectrum β-lactamase (ESBL)-, AmpC- or carbapenemase-producing strains [7, 8, 9, 10, 11]. As a result, the global population structure and lineage diversity of companion-animal-associated *K. pneumoniae* remain poorly understood. Whether these isolates represent distinct host-associated populations or are drawn from the same globally circulating strain pool as human-associated *K. pneumoniae* has yet to be determined.

*K. pneumoniae* is a high-priority AMR pathogen responsible for substantial morbidity and mortality worldwide, particularly when resistant to third-generation cephalosporins (3GCs) or carbapenems [12, 13]. International high-risk clones including ST11, ST15, ST147, ST258 and ST307 have expanded across healthcare systems and are frequently associated with ESBL and carbapenemase genes [14, 15, 16, 17]. Importantly, related *K. pneumoniae* strains and STs have also been identified in companion animals and humans, including evidence of strain sharing within households [8].

Here, a global genome dataset of *K. pneumoniae* from domestic dogs and cats was assembled to define lineage diversity in these companion animals, AMR burden, host structure and relationship to human-associated populations. The distribution of ESBL and carbapenemase determinants, geographic and host-associated MDR patterns, source-associated resistance carriage, dominant STs and high-risk clones was investigated. Companion-animal ST comparisons against a large recent human PathogenWatch population was additionally performed to contextualise whether dog- and cat-associated lineages overlap with globally circulating human *K. pneumoniae* clones.

## Methods

### Genome Dataset Assembly and Metadata Curation

Publicly available *K. pneumoniae* genome records associated with companion animals were identified from the European Nucleotide Archive (ENA), National Center for Biotechnology Information (NCBI) BioSample records and the NCBI Pathogen Detection database, up to May 27, 2026 using host- and source-associated terms for domestic dogs and cats. Metadata fields, including BioSample title, geographic location, collection date, strain name and isolate identifiers, were standardised and merged into Supplementary File 1. Host metadata were standardised to dog or cat using common and scientific names, including *Canis lupus familiaris* and *Felis catus*. Records with human-associated metadata, food or environmental sources, or unresolved companion-animal host assignment were excluded from host-stratified analyses. Geographic metadata were standardised at country level (Supplementary File 1).

### Genome Quality Control and Species Confirmation

Where read-based assembly was required, draft genomes were generated using Shovill v1.1.0 [18] with a target depth of 100, minimum contig length of 500 bp and minimum contig coverage of 5. Assembly quality was assessed from total assembly size, contig count, N50 and longest contig length. Assemblies were excluded if they were empty or outside quality thresholds; total size <4.5 Mb or >7.0 Mb, >500 contigs, N50 <20 kb, or longest contig <50 kb. Species confirmation and downstream genotyping were performed using Kleborate v3.2.4 [19]; population analyses were restricted to isolates confirmed as *K. pneumoniae* from domestic dogs or cats.

### MLST, AMR and Virulence Gene Detection

ST, antimicrobial-resistance determinants, resistance class counts, virulence loci, capsule and O-locus calls were extracted from Kleborate annotations [19]. ESBL and carbapenemase carriage were defined from Kleborate β-lactamase fields, including Bla_ESBL_acquired and Bla_Carb_acquired. Multidrug resistance (MDR) was defined genotypically as resistance determinants spanning at least three antimicrobial classes; extensively drug-resistant isolates were included within the MDR/XDR category. Epidemic/high-risk clone status was assigned to ST11, ST15, ST147, ST307, ST258, ST512, ST101, ST14, ST23, ST37 and ST392.

### Geographic, Host and Source-Stratified Analyses

Country- and region-level summaries were generated for isolate counts, ST composition, MDR/XDR burden, ESBL carriage and carbapenemase carriage. Isolation sources were grouped as infection-associated, colonisation/screening-associated or unknown using local and NCBI metadata. Infection-associated terms included urine, bladder, blood, wound, abscess, eye, lung, clinical, diseased and infection; colonisation/screening terms included faeces/feces, rectal, stool, perianal and screening.

### Lineage Diversity and Statistical Analysis

ST richness, Shannon diversity and Simpson diversity were calculated for host, country and regional strata. Dog and cat sample sizes were unequal, therefore, dog ST richness was rarefied to the cat sample size using Monte Carlo resampling. Host-associated differences in ST composition were assessed using PERMANOVA based on categorical ST dissimilarity. Fisher exact tests were used for binary host comparisons, chi-square tests for regional associations and Spearman correlations for country-level relationships between MDR burden and lineage diversity. Logistic regression was used to estimate the association between cat origin and MDR/XDR carriage, with sequential adjustment for epidemic-clone status, country group, collection year and source category. Statistical analyses were performed in Python v3.11.15 using pandas v3.0.2, NumPy v2.4.3, SciPy v1.17.1 and Statsmodels v0.14.6. Benjamini-Hochberg correction was applied where multiple testing was performed.

### ST147 Comparator Genome Selection and Recombination-Filtered Phylogenomics

ST147 phylogenomics used companion-animal ST147 genomes together with contextual human and animal comparator genomes to construct a recombination filtered phylogeny for feline isolates detected in 2025-6. A lineage-matched internal reference was selected using Mash v2.3 [20] by identifying the genome with the lowest mean pairwise distance. Core-genome alignments were generated with Parsnp v2.1.5 and converted with harvesttools v1.3, both from the Harvest suite [21]. Recombinant regions were identified and masked using Gubbins v3.4.2 [22]. Maximum-likelihood phylogenies were inferred from recombination-filtered polymorphic sites using IQ-TREE v3.1.1 under a GTR+G model with 1,000 ultrafast bootstrap replicates [23]. Pairwise recombination-filtered SNP distances were calculated using snp-dists v1.2.0.

### Plasmid Replicon Profiling

Plasmid replicon profiles were assigned using BLASTN v2.12.0+ against a local PlasmidFinder replicon database [24]. Replicons were considered present using thresholds of ≥95% nucleotide identity and ≥60% replicon coverage. Replicon profiles were interpreted alongside core-genome phylogeny, host category, collection year and AMR genotype to assess accessory-genome conservation within the ST147 lineage.

### Construction of a Global Human *K. pneumoniae* Comparator Dataset

A contemporaneous *K. pneumoniae* human derived comparator dataset was generated to compare ST similarity with the companion animal dataset. Public PathogenWatch records assigned to *K. pneumoniae* were queried through the PathogenWatch application programming interface (API). For each record, isolate identifier, ST, collection date and geographic metadata were retained where available. BioSample accessions from PathogenWatch were then queried against the ENA portal API to retrieve sample-level host and isolation-source metadata. The resulting PathogenWatch and ENA metadata tables were merged by BioSample accession, after which records were filtered to retain only human-associated isolates, defined by host/source annotations labelled *Homo sapiens*, human or patient. To provide recent *K. pneumoniae* isolates for comparison, records were retained only if the collection year was >=2019 and a valid multi-locus sequence type (MLST) assignment was available. The human comparator dataset (available on Github, https://github.com/StephenFordham/-Human_K.-pneumoniae_Comparator_Dataset_2019_onwards_SF/blob/main/humandataset.csv) comprises 38,106 human-associated *K. pneumoniae* isolates spanning 1,661 STs and was used for human-animal ST distribution comparisons.

## Results

### H2ost-shared dominant lineages show distinct dog and cat enrichment

Across 712 domestic dog- and cat-derived *K. pneumoniae* isolates, 263 STs were detected, indicating substantial lineage diversity in the companion-animal population (Supplementary File 1). Despite this diversity, the ten most frequent STs accounted for 326/712 isolates (45.8%), comprising ST307 (*n*=84), ST11 (*n*=75), ST15 (*n*=51), ST147 (*n*=40), ST37 (*n*=25), ST45 (*n*=12), ST17 (*n*=10), ST392 (*n*=10), ST1 (*n*=10), and ST16 (*n*=9). All ten STs were detected in both dogs and cats, supporting host sharing rather than host restriction (Figure 1). However, their contribution differed by host: top-10 STs accounted for 236/567 dog isolates (41.6%), but 90/145 cat isolates (62.1%), indicating that cat-derived isolates were more concentrated within dominant lineages. The three most common STs, ST307, ST11, and ST15, represented 210/712 isolates (29.5%) and were predominantly dog-associated, although each was also recovered from cats. In contrast, *K. pneumoniae* ST147 was the only top 10-ST detected more often in cats (24 versus 16), reinforcing its cat-enriched structure within an otherwise dog-heavy population (Figure 1). Temporal metadata showed repeated detection of several major STs across recent years, with ST307 and ST11 represented from 2020 to 2026, ST15 from 2020 to 2025, and ST147 in 2023, 2025, and 2026 (Figure 1). Overall, companion animal-associated *K. pneumoniae* is highly diverse but includes a concentrated subset of recurrent, host-shared dominant lineages. The most common STs are dog-enriched, however ST147 represents a cat-enriched high-risk clone.

**Figure 1.**
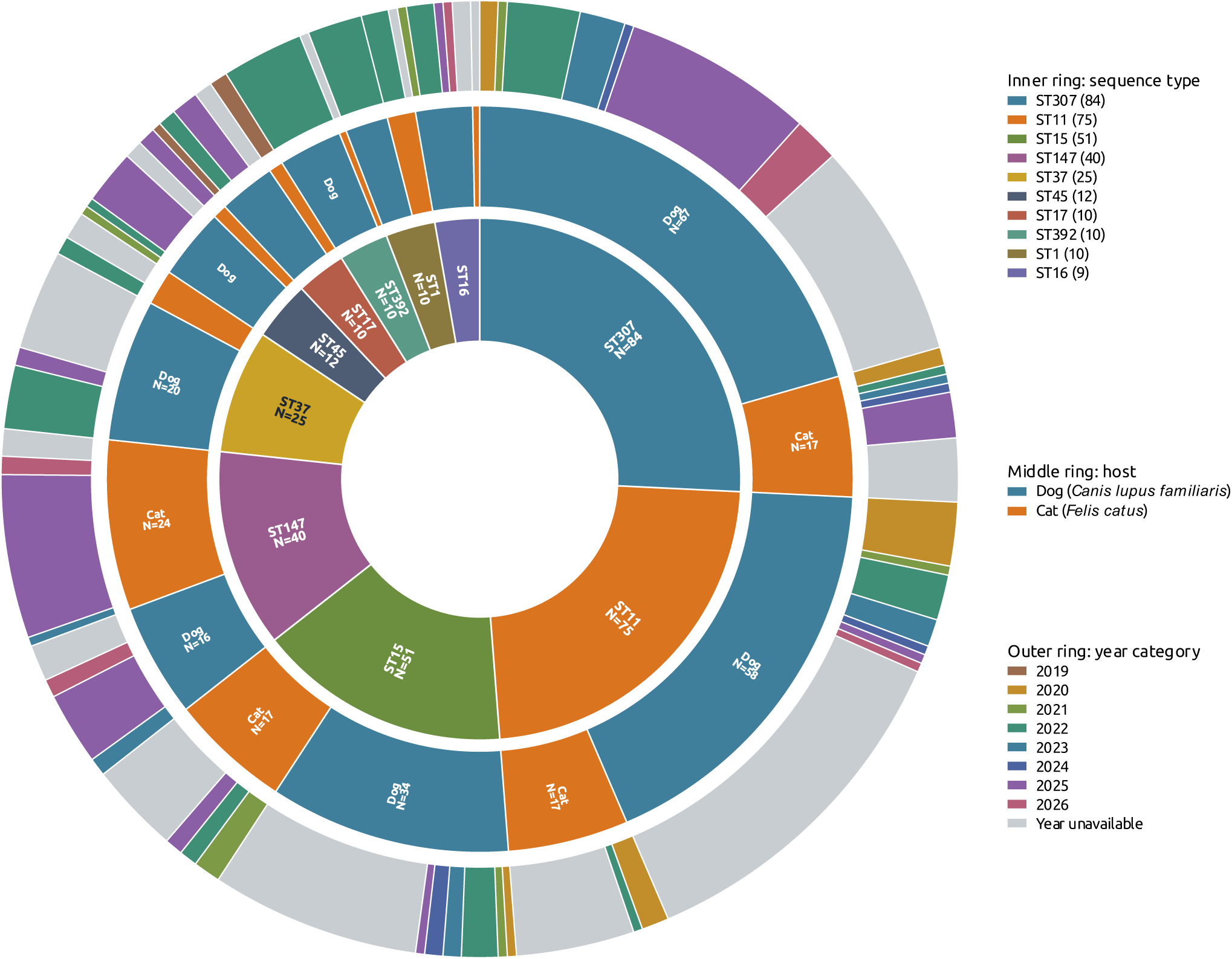
Host and temporal structure of dominant companion animal-associated *K. pneumoniae* lineages. Circular ring count plot shows the ten most frequent STs among domestic dog- and cat-derived *K. pneumoniae*. The inner ring shows the relative abundance of each ST, the middle ring partitions each ST by host species, and the outer ring shows collection-year categories, including isolates without available year metadata. The dominant lineages were shared across host species but showed clear differences in host representation. ST307, ST11, and ST15 were the most abundant STs overall and were predominantly dog-associated, whereas ST147 showed a comparatively stronger contribution from cat-derived isolates while also including dog-derived genomes. The year-stratified outer ring indicates that several major STs were detected across multiple collection years, supporting recurrent recovery of these lineages from companion animals rather than isolated single-time-point observations. This structure highlights the coexistence of broadly distributed dog-associated lineages and cat-enriched high-risk lineages within the companion-animal *K. pneumoniae* population.

### Comparison of Companion-Animal and Human *Klebsiella pneumoniae* Sequence Type Distributions

To contextualise companion animal-associated *K. pneumoniae* within global human-associated diversity, the domestic dog and cat dataset was compared with a large human comparator dataset described in the Methods. This comparator included 38,106 human-associated *K. pneumoniae* isolates with ST assignments, collected from 2019 onwards across 91 countries and spanning 1,661 STs. By comparison, the companion-animal dataset comprised 712 isolates spanning 263 MLST profiles.ST sharing was extensive: 187/263 companion-animal MLST profiles (71.1%) were also present in the human comparator, and 621/712 companion-animal isolates (87.2%) belonged to STs profiles detected among human-associated *K. pneumoniae*. All 10 most frequent companion-animal STs were present in the human dataset, 19/20 of the top companion-animal STs were detected in humans, and 10/20 were also among the 20 most frequent human STs. This overlap included several internationally recognised high-risk clones, including ST11, ST15, ST17, ST147, ST307, ST258 and ST101.

The overlap in ST composition was also reflected in lineage rank. Among the ten most frequent companion-animal STs, rank order was positively correlated with that observed in the human *K. pneumoniae* population (Spearman ρ = 0.745, *p* = 0.013; Figure 2a). Across 100 random samples of 10,000 human isolates, companion-animal and human ST frequencies showed a moderate positive correlation among shared STs (mean Spearman ρ = 0.566), with a mean Bray-Curtis dissimilarity of 0.552 and a mean top-20 Jaccard overlap of 0.309. A permutation test confirmed that the observed animal-human ST similarity was substantially greater than expected by chance (observed Bray-Curtis similarity = 0.448 versus null mean = 0.054; *p* = 0.00010, Figure 2b). Among the top ten companion-animal STs, ST147, ST11, ST307, ST45, ST15, ST17 and ST16 all ranked among the ten most common human-associated STs.

High-risk clone burden was also comparable between populations using the high-risk ST list (see method): 262/712 companion-animal isolates (36.8%) and 15,061/38,106 human isolates (39.5%) belonged to these clones (Fisher exact OR = 0.89, *p* = 0.142), indicating that companion animals share a similar proportion of clinically important lineages relative to the human *K. pneumoniae* population.

Temporal metadata additionally supported contemporary overlap for major shared lineages. Among the top 10 companion-animal STs, five had animal records dated between 2019 and 2026 (ST307, ST11, ST15, ST147 and ST17), and all five were also detected among human-associated *K. pneumoniae* in overlapping collection years. ST307 was detected in both datasets in 2022, 2024 and 2025; ST11 in 2023-2025; ST15 in 2023 and 2025; ST147 in 2023, 2025 and 2026; and ST17 in 2019 and 2025. At this population-genetic scale, the extensive sharing of STs and ST-associated profiles between companion animals and humans indicates that domestic dogs and cats harbour lineages drawn from the same broad *K. pneumoniae* strain pool as human-associated isolates, consistent with potential ecological connectivity between host populations.

**Figure 2.**
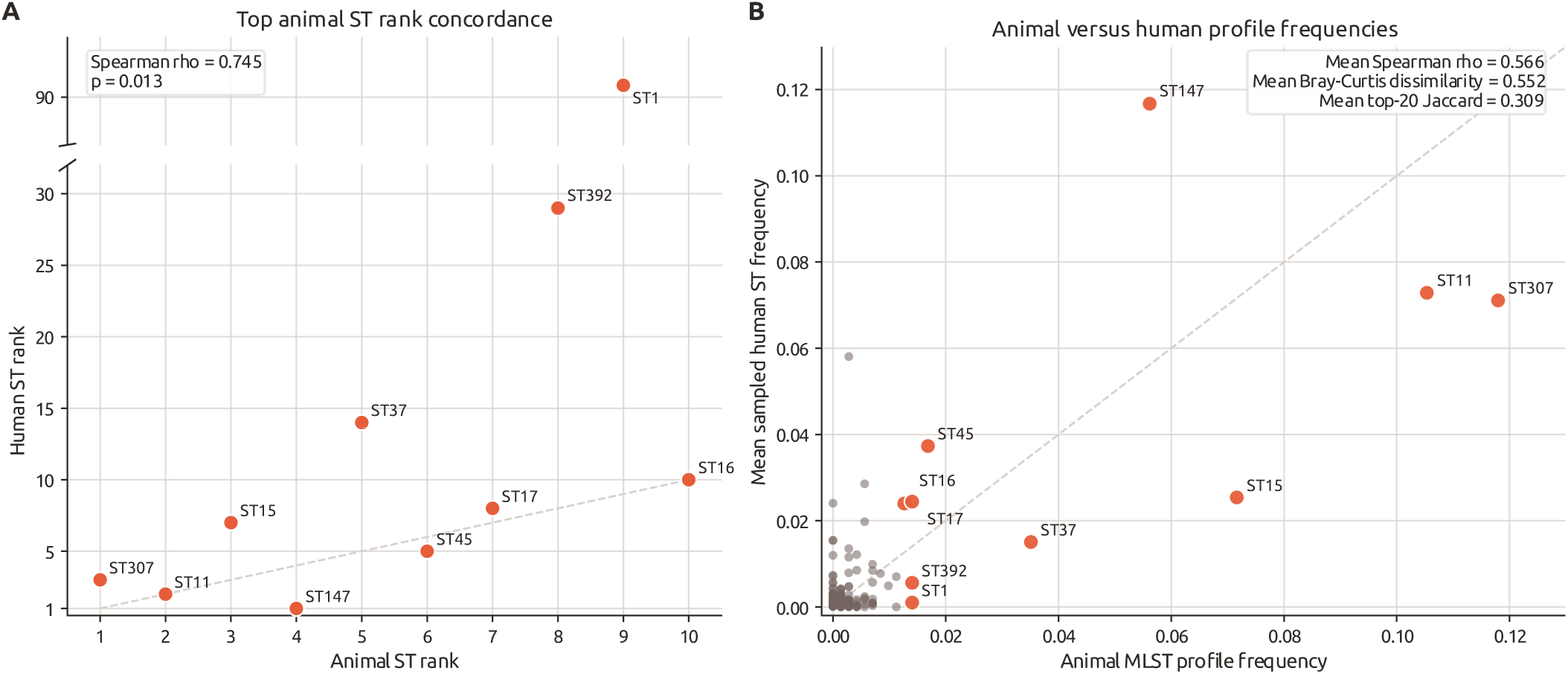
Concordance between companion animal-associated and human-associated *K. pneumoniae* sequence type distributions. **(A)** Rank concordance between the ten most frequent domestic dog/cat-associated *K. pneumoniae* STs and their corresponding ranks in the human comparator dataset. ST1 is shown using a broken y-axis because it was common in the companion-animal dataset but comparatively uncommon among human-associated isolates. Rank concordance was significant across the top ten companion-animal STs (Spearman ρ = 0.745, p = 0.013). **(B)** Scatter plot comparing companion-animal MLST-profile frequency with mean sampled human ST frequency. Human frequencies were estimated from 100 random samples of 10,000 human-associated *K. pneumoniae* isolates collected from 2019 onwards. The ten most frequent companion-animal STs are highlighted in red; all other MLST profiles are shown in brown. Across sampling iterations, companion-animal and human distributions showed moderate similarity among shared profiles (mean Spearman ρ = 0.566), with incomplete but non-random overlap in overall frequency structure (mean Bray-Curtis dissimilarity = 0.552; mean top-20 Jaccard index = 0.309). Together, these analyses show that dominant companion-animal *K. pneumoniae* lineages overlap substantially with globally circulating human-associated STs, while retaining differences in relative abundance.

### Global dissemination of ESBL and carbapenemase resistance determinants in companion animal-associated *K. pneumoniae*

To determine whether clinically important β-lactam resistance determinants are disseminated among companion animal-associated *K. pneumoniae*, we examined ESBL and carbapenemase gene carriage among *K. pneumoniae* isolates recovered from domestic dogs and cats with available country metadata (*n*=706). ESBL genes were common, being detected in 303/706 isolates (42.9%) and spanning 23 countries across four major geographic regions. Carbapenemase genes were less frequent but still widely distributed, occurring in 98/706 isolates (13.9%) across eight countries and three regions. In total, 345/706 isolates (48.9%) carried either an ESBL or carbapenemase gene, indicating that high-priority β-lactam resistance determinants are widespread in the companion animal collection (Supplementary File 1).

The burden of ESBL/carbapenemase carriage varied substantially by geography. The largest absolute numbers of ESBL/carbapenemase-positive isolates were identified in the USA (96/231, 41.6%), France (71/108, 65.7%), China (65/124, 52.4%), South Korea (26/29, 89.7%), Switzerland (12/14, 85.7%), Brazil (11/12, 91.7%), Germany (10/52, 19.2%), and Spain (10/14, 71.4%). At regional scale, Asia had the highest proportion of ESBL/carbapenemase-positive isolates (95/157, 60.5%), followed by Europe (135/281, 48.0%), the Americas (112/259, 43.2%), and Africa (3/8, 37.5%); no ESBL or carbapenemase genes were detected in the single isolate from Oceania. Geographic region was significantly associated with ESBL/carbapenemase carriage (chi-square (χ^2^) =13.24, df = 4, *p* = 0.0102), although this finding should be interpreted in the context of uneven sampling intensity across countries and regions.

The dissemination of these resistance determinants was strongly linked to dominant *K. pneumoniae* lineages. ESBL/carbapenemase-positive isolates were concentrated in several internationally recognised high-risk STs, including ST307 (*n*=80 carriers across 13 countries and three regions), ST15 (*n*=49 across nine countries and three regions), ST11 (*n*=42 across nine countries and three regions), and ST147 (*n*=33 across five countries and three regions). This pattern indicates that global dissemination is associated with the expansion and geographic spread of successful resistant lineages.

*bla*_CTX-M-15_ was the dominant ESBL determinant identified, detected in 231 isolates across 22 countries, consistent with broad international dissemination of this ESBL determinant in domestic dog/cat animal-associated *K. pneumoniae*. Other ESBLs were less frequent but included *bla*_CTX-M-55_ (*n*=19), *bla*_CTX-M-27_ (*n*=16), *bla*_CTX-M-14_ (*n*=13), and *bla*_CTX-M-3_ (*n*=10). Among carbapenemases, *bla*_NDM-5_ was most common (*n*=48 isolates across four countries), followed by *bla*_OXA-48_ (*n*=31 across four countries), *bla*_KPC-3_ (*n*=6), *bla*_NDM-7_ (*n*=6), *bla*_OXA-181_ (*n*=3), *bla*_OXA-232_ (*n*=3), and *bla*_KPC-2_ (*n*=2). Together, these results demonstrate that ESBL-producing companion animal-associated *K. pneumoniae* are globally disseminated, while carbapenemase-producing isolates, although less prevalent, are also distributed internationally and are concentrated within high-risk lineages. These findings support companion animals as a reservoir for globally important β-lactam resistance determinants and highlight the need to include domestic dogs and cats in genomic surveillance frameworks for antimicrobial-resistant *K. pneumoniae*.

As ESBL and carbapenemase genes represent only part of the wider resistance burden, we next assessed MDR/XDR structure across countries and regions to determine whether geographic variation reflected overall resistance load, lineage composition, or sampling intensity.

### Geographic variation in MDR burden reflects regional lineage structure and sampling intensity

Geographic MDR burden and lineage diversity were assessed using *K. pneumoniae* dog/cat isolates with country metadata (*n*=706). MDR was defined as carriage of resistance genes across >=3 AMR classes, including XDR isolates. The largest MDR burdens were observed in France/Guadeloupe (*n*=106/117 MDR; 90.6%), the USA (*n*=110/231; 47.6%), China (*n*=82/124; 66.1%), South Korea (*n*=27/29; 93.1%), and Germany (*n*=16/52; 30.8%) (Figure 3a). Among countries represented by at least 10 isolates, the highest MDR proportions were observed in Switzerland, Brazil, and Spain (all 100%), followed by South Korea, Portugal, and France/Guadeloupe.

Regional clone structure revealed that Europe had 51.2% of isolates assigned to recognised epidemic clones, dominated by ST11, ST15, and ST307. ST11 was significantly enriched in Europe (OR=9.95, BH-adjusted *q* = 6.15 × 10^−15^). Asia showed significant enrichment of ST392 (OR = 14.7, *q* = 0.00169) and ST37 (OR = 4.04, *q* = 0.00596), while the Americas showed significant enrichment of ST147 (OR = 6.62, *q* =2.14 × 10^−6^). Lineage diversity was highest in the Americas (140 STs; Shannon H=4.278), followed by Europe (105 STs; H=3.568) and Asia (71 STs; H=3.752).

At country level, the USA had the greatest observed lineage diversity. Country-level lineage diversity was negatively correlated with MDR proportion; Shannon diversity versus MDR proportion gave Spearman ρ = -0.676 (*p* = 0.0158). Similarly, Simpson diversity versus MDR proportion gave ρ = - 0.725 (*p* = 0.00759), and ST richness versus MDR proportion gave ρ = -0.596 (*p* = 0.0408). These results suggest that countries dominated by fewer lineages have higher MDR proportions, whereas absolute MDR counts were more strongly influenced by sampling intensity.

**Figure 3.**
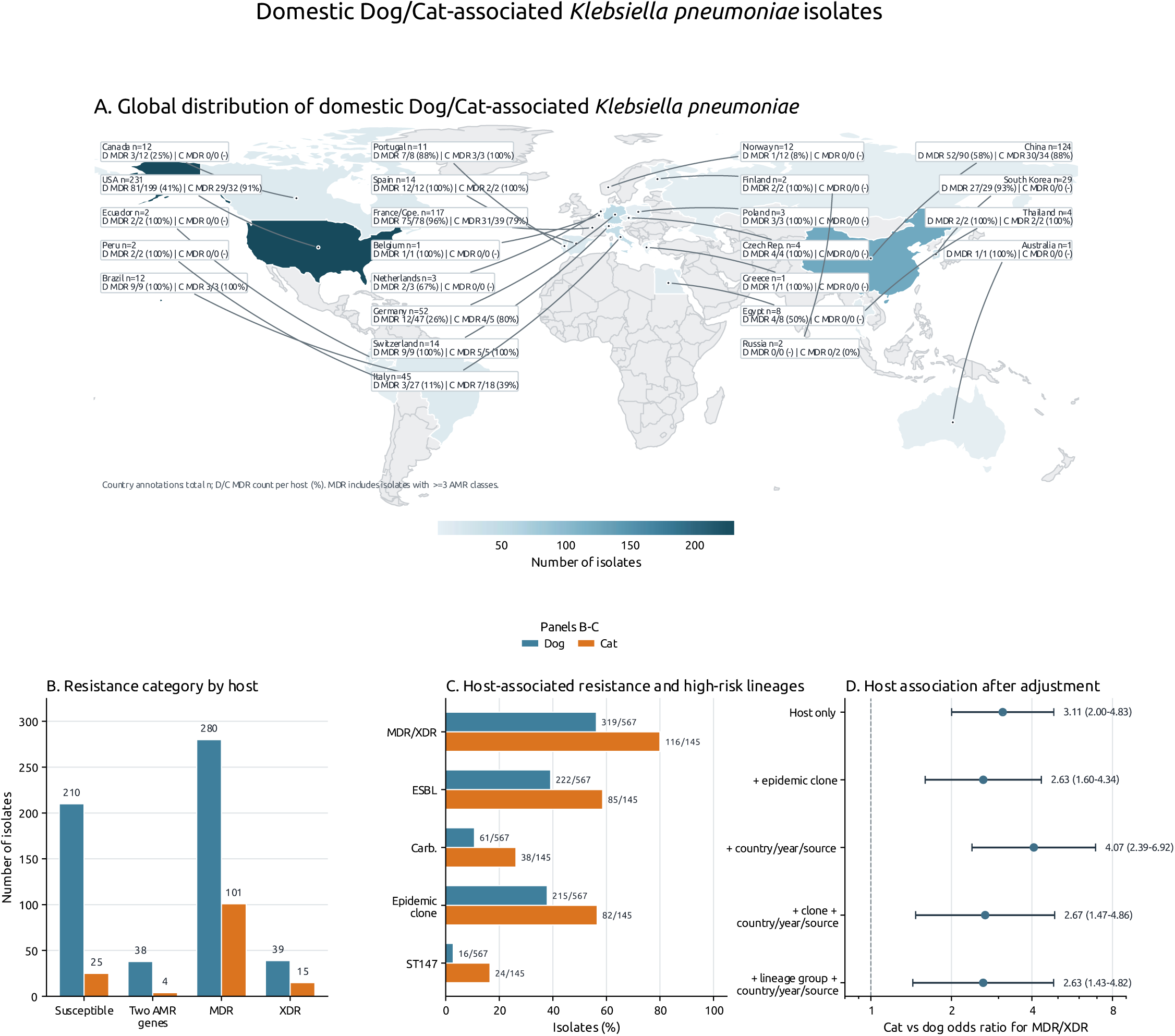
Global distribution, resistance burden, and host-associated risk structure of domestic dog- and cat-associated *K. pneumoniae*. 712 *K. pneumoniae* isolates recovered from domestic dogs (*n* = 567) and cats (*n* = 145). **A.** Country-level distribution of isolates, with shading indicating the number of isolates per country. Country annotations show total isolate counts and host-stratified MDR burden for dogs and cats. **B.** Genotypic resistance categories among dog-and cat-derived isolates, classified as susceptible, carrying two antimicrobial resistance genes, multidrug-resistant (MDR), or extensively drug-resistant (XDR). **C.** Host-stratified prevalence of key resistance and lineage features, including MDR/XDR genotype, ESBL genes, carbapenemase genes, epidemic clone membership, and ST147. Cat-derived isolates showed a higher proportion of MDR/XDR genotypes than dog-derived isolates (116/145, 80.0% vs 319/567, 56.3%), and were also more frequently associated with ESBL genes, carbapenemase genes, epidemic clone membership, and ST147. **D.** Logistic regression estimates for the association between cat origin and MDR/XDR carriage. Points represent cat-versus-dog odds ratios and horizontal lines show 95% confidence intervals. The association persisted after adjustment for epidemic clone status, country, year, isolation source, and broader lineage structure, supporting enrichment of MDR/XDR among cat-derived isolates while highlighting the contribution of clonal and epidemiological structure. MDR was defined as resistance determinants spanning three or more antimicrobial classes.

### Domestic dogs harbour broader and more even *K. pneumoniae* ST diversity than cats

The geographic analyses indicated that resistance burden was closely linked to lineage structure; therefore, we next compared ST diversity directly between dog- and cat-derived isolates to determine whether host species differed in the breadth and evenness of their *K. pneumoniae* populations. Among *K. pneumoniae* isolates with ST assignment calls belonging to either domestic dogs or cats (*n*=712; dogs *n*=567, cats *n*=145), dogs harboured substantially more diverse ST lineages than cats. Dogs had higher observed ST richness (240 STs vs 51 in cats), higher Shannon diversity (H=4.569 vs 3.187; permutation test, delta H=1.382, *p* = 0.0001, 9,999 permutations), and lower lineage dominance/higher Simpson diversity (1-D=0.967 vs 0.924; permutation *p* = 0.0004). As sampling was unequal, richness was rarefied to the cat sample size (*n*=145). Here, dogs still had higher expected ST richness (84.08 STs; Monte Carlo 95% interval 75-93) than cats (51 STs), with a rarefied richness difference of 33.08 STs (*p* = 0.0001). PERMANOVA based on categorical ST dissimilarity showed that host species was also associated with a statistically significant difference in lineage composition (pseudo-F=3.357, R^2^=0.0047, *p* = 0.0001, 9,999 permutations), although the low R^2^ indicates that host explains only a small fraction of total ST compositional variation. Overall, the data support that dog-derived isolates contain a broader and more even set of *K. pneumoniae* ST lineages than cat-derived isolates.

### Higher MDR/XDR prevalence among cat-derived isolates is partly explained by high-risk lineage structure

Cat-derived *K. pneumoniae* isolates showed a higher MDR/XDR prevalence than dog-derived isolates (116/145, 80.0% versus 319/567, 56.3%; Fisher exact OR = 3.11, 95% CI 2.00-4.83; p=7.29 × 10^−8^). Cat isolates also more frequently carried ESBL genes (58.6% versus 39.2%; OR=2.20, 95% CI 1.52-3.19; p=3.23 × 10^−5^) and carbapenemase genes (26.2% versus 10.8%; OR = 2.95, 95% CI 1.87-4.65; *p*=6.57 × 10^−6^) (Figure 3c). This indicates that the higher MDR/XDR burden in cats coincided with enrichment for clinically important β-lactam resistance determinants.

This host-associated pattern was partly explained by lineage structure. Cat isolates were more likely than dog isolates to belong to recognised epidemic/high-risk clones (ST11, ST15, ST147, ST307, ST258, ST512, ST101, ST14, ST23, ST37 and ST392) (56.6% versus 37.9%; OR = 2.13, 95% CI 1.47-3.08; *p* = 6.78 × 10^−5^). ST147 was also strongly over-represented in cats (16.6% versus 2.8%; OR=6.83, 95% CI 3.52-13.25; *p*=1.62 × 10^−8^) (Figure 3c). In logistic regression, cat origin was associated with MDR/XDR carriage in a host-only model (OR=3.11, 95% CI 2.00-4.83; *p*= 4.19 × 10^−7^) (Figure 3d). Adjustment for epidemic/high-risk clone status reduced, but did not remove, this association (OR = 2.63, 95% CI 1.60-4.34; *p* = 1.51 × 10^−4^), while epidemic-clone carriage itself was strongly associated with MDR/XDR (OR = 18.01, 95% CI 11.23-28.87; *p* = 3.51 × 10^−33^).

Country, year, and source did not explain away the cat-associated MDR/XDR signal. Adjustment for country, collection year, and source category gave a cat-associated OR of 4.07 (95% CI 2.39-6.92; *p* = 2.37 × 10^−7^), while a model additionally including epidemic-clone status gave an OR of 2.67 (95% CI 1.47-4.86; *p*=0.00128) (Figure 3d). A reduced lineage model separating ST147, other epidemic clones, and non-epidemic lineages, while also adjusting for country, year, and source, gave a similar cat effect (OR=2.63, 95% CI 1.43-4.82; *p* = 0.00178). As full ST fixed-effect models were unstable in this dataset due to rare STs and near-complete MDR/XDR in several lineages, ST-aware inference was also assessed using a within-ST permutation test restricted to STs present in both hosts. This retained a host-associated difference in MDR/XDR carriage (28 ST strata; 399 isolates; observed cat-minus-dog difference=7.3 percentage points; 99,999 permutations, *p*=0.0197).

Overall, cat-derived isolates had higher MDR/XDR prevalence than dog-derived isolates, and this association persisted after adjustment for measured country, year, source, and broad lineage structure. However, the effect was attenuated after accounting for epidemic/high-risk clones, indicating that clonal composition contributes to the host difference. These results should therefore be interpreted as an epidemiological association rather than evidence of host-driven causation. Higher MDR/XDR prevalence among cat-derived isolates may also reflect a combination of high-risk clone expansion, sampling framework, geography, veterinary healthcare exposure, antimicrobial use history, and host-associated epidemiology, rather than cats being more likely to harbour MDR *K. pneumoniae*.

### MDR *K. pneumoniae* are not restricted to infection-associated samples

MDR carriage was not confined to infection-associated isolates. MDR was detected in 82/132 colonisation/screening-associated isolates (62.1%) compared with 193/368 infection-associated isolates (52.4%). Although the observed proportion was higher among colonisation/screening-associated isolates, this difference did not reach statistical significance in the unadjusted comparison (Fisher exact test, OR = 0.67 for infection-associated versus colonisation/screening-associated isolates, 95% CI 0.45-1.01, *p* = 0.0663).

After adjustment for host species, the source-associated difference was further attenuated and remained non-significant (adjusted OR = 0.82, 95% CI 0.53-1.26, *p* = 0.354), indicating that host composition partly explains the observed unadjusted pattern. These results remain important because they indicate MDR *K. pneumoniae* are frequently recovered from colonisation or screening-associated sources as well as infection-associated sources. This supports the potential role of domestic dogs and cats as important reservoirs of MDR lineages beyond clinically diseased animals.

### A host-spanning MDR ST147 sub-lineage links companion-animal and human-associated *K. pneumoniae*

As ST147 was both cat-enriched and strongly associated with clinically important resistance determinants, with strains recently detected in 2025, and 2026, we performed focused phylogenomic analysis to determine whether companion animal ST147 represented a host-restricted expansion or part of a broader host-spanning lineage.

Recombination-aware phylogenomic analysis placed the companion-animal ST147 *K. pneumoniae* isolates within a structured and heterogeneous ST147 population, rather than within a single animal-restricted lineage. A cat-enriched sub-lineage was identified. This lineage, however, was not exclusively feline: dog-derived isolates, additional animal-context isolates, and closely related human comparator genomes were embedded within or immediately adjacent to the same genomic background. In contrast, other ST147 cat and dog genomes were positioned in more distant parts of the tree, indicating multiple introductions or acquisitions of ST147 by companion animals rather than expansion of a single animal-associated clone.

The feline-dominant lineage showed limited core-genome diversity. Among the 25 cat and dog-derived isolates in this lineage, pairwise recombination-filtered distances ranged from 0 to 34 SNPs, with a median of 6 SNPs. The 17 USA-NY feline isolates formed an especially tight cluster, differing by 0 to 28 SNPs, with a median of only 3 SNPs. Using the USA-NY cat isolate 144D-D (SAMN48918375) as a fixed feline-clade comparator, several cat isolates and both USA-NY dog isolates were 0 SNPs from this genome, while most remaining USA-NY cat isolates were within 1 to 5 SNPs (Figure 4). These distances are consistent with recent diversification of a highly related ST147 sub-lineage circulating across companion-animal hosts.

Human comparator genomes were closely associated with the animal-associated cluster. The nearest human genome to each animal isolate, within the feline dominated sub-lineage was separated by only 2 to 9 recombination-filtered SNPs; across all animal-to-human comparisons within the NY feline-dominated lineage, distances ranged from 2 to 37 SNPs, with a median of 22 SNPs. Several USA-NY cat isolates were 3 to 4 SNPs from human isolate 2025HL-00779, and dog-derived isolate 45618-23 was 2 SNPs from human isolate 2024DK-00378. These data support a host-spanning ST147 lineage with very recent shared ancestry between animal- and human-associated genomes. However, in the absence of epidemiological linkage, these phylogenetic relationships should be interpreted as evidence of a shared circulating background rather than proof of direct transmission. The distinctiveness of the focal lineage was further supported by comparison with ST147 genomes outside this sub-lineage: distances from the animal isolates within the cat-dominated sub-lineage to external ST147 comparators were substantially larger, ranging from 93 to 298 SNPs, with a median of 130 SNPs (Figure 4).

The host distribution, resistance profile and accessory-genome structure together define a cat-enriched, MDR ST147 sub-lineage that is not host restricted. Within the compact animal-associated lineage, all 25 cat- and dog-derived isolates carried the same core resistance variants, comprising the carbapenemase gene *bla*_NDM-5_, the quinolone-resistance gene *qnrB1* and the trimethoprim-resistance gene *dfrA14*, indicating a conserved MDR genotype across the companion-animal component of the clade. However, the phylogeny does not support a strictly feline lineage: dog-derived isolates, animal-context isolates and closely related human comparator genomes were interspersed within, or positioned immediately adjacent to, the same genomic background (Figure 4). These results indicate a recent expansion of a highly related MDR ST147 population enriched among cats and extending across companion-animal and human-associated genomes. This is supported by the low recombination-filtered SNP distances within the feline-dominated cluster and the close proximity of human comparator isolates.

Plasmid replicon profiling further showed that this lineage sits within a broader ST147 accessory-genome background dominated by recurrent Inc groups. Across the complete set of 57 ST147 genomes analysed by recombination-filtered core-genome phylogeny and plasmid replicon profiling, the most frequent replicon groups were IncFIB(K), IncFII(K) and IncX3, detected in 46/57, 45/57 and 41/57 genomes, respectively (Figure 4). These replicons were present across human, cat, dog-derived and animal-context isolates, and were retained across the feline-dominated clade rather than being confined to a single host category. Among dated recurrent-replicon-positive genomes, all three major replicons were already present in the 2022 human isolate 2022HL-01876 (SAMN31854197), were subsequently detected in the 2023 dog-derived isolate 45618-23 (SAMN40717015), and persisted through the 2024-2026 expansion that included human comparators, the USA-NY cat cluster, and later dog-derived isolates sampled in 2025 and 2026. This temporal pattern supports persistence of a conserved IncFIB(K)/IncFII(K)/IncX3 plasmid-replicon architecture within the ST147 population before and during the observed companion-animal expansion.

In contrast to these dominant recurring replicons, additional Inc groups were uncommon and generally restricted to one or two genomes, consistent with episodic plasmid gain, loss or replacement against a stable accessory-genome backbone. The combined core-genome and plasmid-replicon data therefore suggest that the feline-dominated ST147 clade did not arise through acquisition of a wholly distinct plasmid profile after establishment in cats. Instead, it appears to represent a recently expanded, host-spanning MDR ST147 lineage carrying a conserved plasmid-replicon repertoire that was already present in closely related human- and dog-associated genomes. The convergence of low core-genome SNP distances, conserved MDR determinants and recurrent plasmid replicons highlights the value of integrated human and companion-animal genomic surveillance for detecting high-risk *K. pneumoniae* lineages at the human-animal interface.

**Figure 4.**
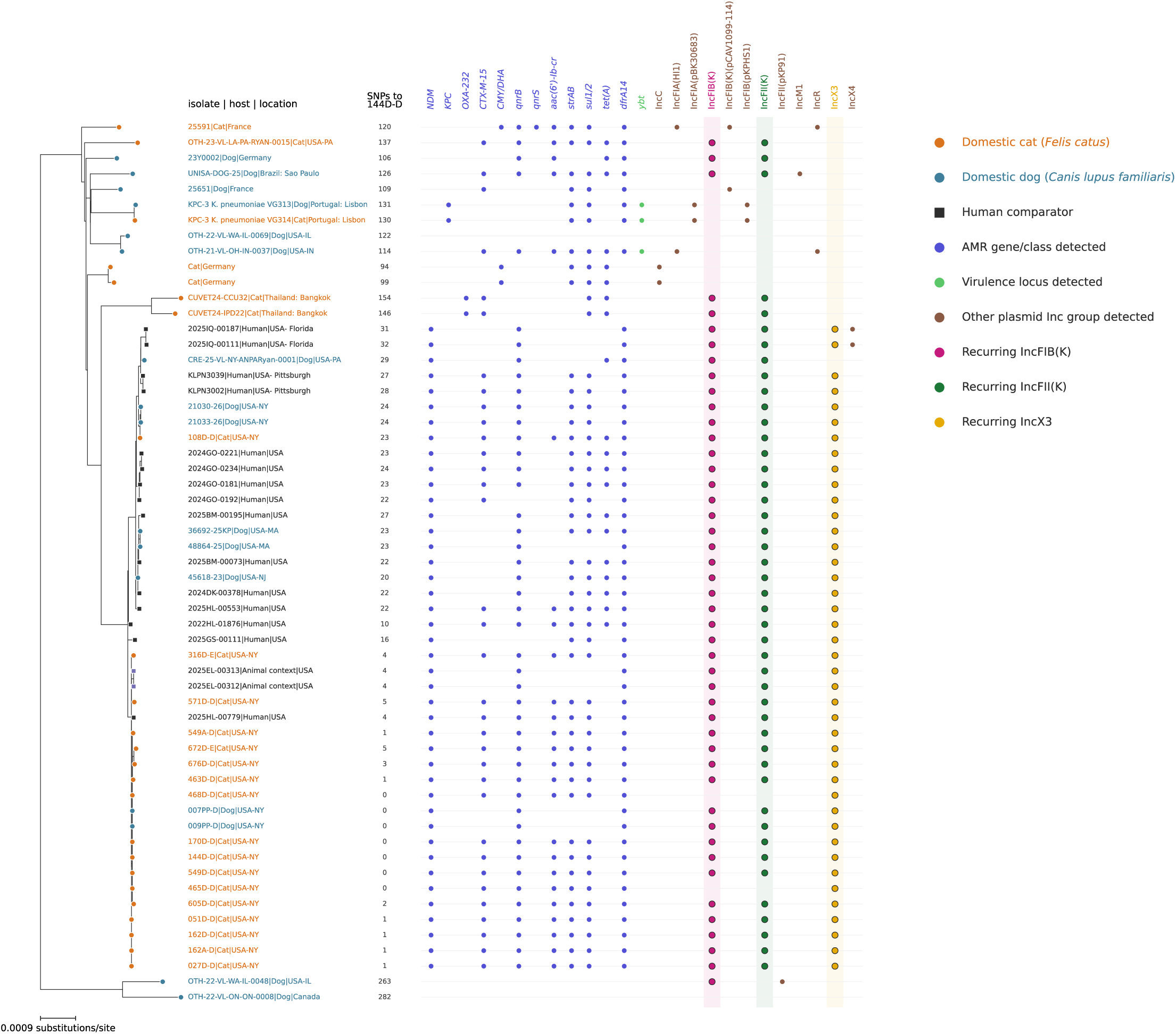
Recombination-filtered core-genome phylogeny of ST147 *K. pneumoniae* isolates from companion animals and contextual human genomes. The tree includes 57 genomes comprising all isolates from the NCBI Pathogen Detection PDS000180176.25 SNP cluster together with additional ST147 cat and dog isolates identified in the companion-animal dataset. Assemblies were aligned with Parsnp using the genome with the lowest mean Mash distance as reference, generating a 4,434,714 bp core-genome alignment. Recombinant regions were masked with Gubbins, leaving 1,054 recombination-filtered polymorphic sites for maximum-likelihood phylogenetic inference with IQ-TREE. Tip colours indicate host category: domestic cat (orange), domestic dog (blue), human comparators (black), and other animal comparators (purple). The SNP column shows recombination-filtered core SNP distances to the fixed USA-NY cat comparator 144D-D (SAMN48918375), selected as a representative of the feline-dominated USA-NY clade. Filled circles indicate presence of selected antimicrobial-resistance determinants, virulence loci, or plasmid incompatibility (Inc) groups. Recurrent IncFIB(K), IncFII(K), and IncX3 groups are highlighted. AMR and virulence calls were derived from Kleborate v3.2.4; plasmid Inc groups were assigned by BLAST against a local PlasmidFinder replicon database using >=95% identity and >=60% replicon coverage thresholds. The phylogeny shows that several canine ST147 isolates fall within or close to the cat-dominated clade, with human comparator genomes included to provide broader genomic context.

## Discussion

Companion animal-associated *K. pneumoniae* are not confined to a distinct veterinary lineage pool but instead include many of the same ST backgrounds that dominate human-associated populations. Across 712 dog- and cat-derived *K. pneumoniae* isolates, 263 MLST profiles were identified, including globally disseminated high-risk clones such as ST307, ST11, ST15 and ST147, lineages associated with international AMR dissemination and high clinical concern [13, 16]. *K. pneumoniae* is a major contributor to the global burden of bacterial AMR and a highly plastic opportunistic pathogen with a well-recognised capacity to acquire and disseminate antimicrobial-resistance determinants across ecological niches [12, 25]. The recovery of these lineages from companion animals provides support that dogs and cats form part of the broader One Health ecology of *K. pneumoniae*, rather than representing isolated hosts for unrelated bacterial populations. This is consistent with increasing evidence that antimicrobial-resistant organisms of human clinical relevance occur in companion animals, households and veterinary settings [10].

The ST sharing observed, provides evidence of population overlap and ecological connectivity. Public genomic datasets, however, also carry sampling biases related to geography, sequencing capacity, outbreak investigation and preferential submission of resistant isolates, which may influence apparent lineage frequencies [26]. However, the scale and consistency of the overlap observed, together with the enrichment of recognised high-risk clones, indicates that companion animals harbour *K. pneumoniae* lineages drawn from the same wider strain pool as human-associated isolates.

The similar MDR rates in infection-associated and colonisation/screening-associated *K. pneumoniae* suggest resistant strains are not found only in animals with clinical disease but can also be carried by animals without obvious infection. This is epidemiologically important because colonisation can provide a clinically silent reservoir from which later infection may arise. In humans, gastrointestinal carriage of *K. pneumoniae* is increasingly recognised as a precursor state for infection, with studies showing that colonising strains can subsequently cause clinical disease and that resistant *K. pneumoniae* may persist unnoticed in healthcare-associated populations [27]. Comparable patterns have been reported in companion animals, where longitudinal studies have shown recurrent or persistent faecal carriage of ESBL/AmpC-producing Enterobacterales in dogs, including post-discharge carriage with associated household environmental contamination [28, 29]. Household studies have also identified carriage, acquisition and co-carriage of ESBL-producing Enterobacterales among dogs, cats and cohabiting humans [30]. Together, these findings support colonisation-positive companion-animal isolates as epidemiologically relevant rather than incidental findings.

The higher MDR prevalence among cat-derived *K. pneumoniae* isolates was partly explained by differences in lineage structure. Cats were significantly enriched for recognised epidemic/high-risk clones compared with dogs, 56.6% versus 37.9%, particularly ST147 (16.6% versus 2.8%), and epidemic-clone carriage was itself strongly associated with MDR status. Adjustment for epidemic-clone membership reduced the cat-associated MDR OR from 3.11 to 2.63, indicating that clonal composition contributed substantially to the observed host-associated difference. This is further supported by the lower lineage diversity observed in cats, which harboured fewer STs, 51 versus 240 in dogs. These STs were also more concentrated within dominant lineages. Importantly, these dominant feline-associated lineages were highly MDR-associated, including all ST147 isolates and 57/58 isolates belonging to other epidemic clones.

Although clonal composition was the strongest measurable explanation for higher MDR prevalence in cats, the factors shaping this lineage structure remain uncertain. Veterinary healthcare exposure, recurrent urinary disease, prior antimicrobial treatment and repeated exposure to similar antimicrobial classes are recognised drivers of MDR Enterobacterales in companion animals [31, 32], but these metadata were unavailable for analysis here. Cat-derived isolates from China were enriched for urinary sources, consistent with previous reports linking feline *K. pneumoniae* infection to urinary tract disease [33], although this pattern was not observed globally. Household transmission and prolonged asymptomatic carriage are also plausible contributors, supported by reports of strain sharing between companion animals and cohabiting humans [8, 34], but these mechanisms cannot be tested without paired owner-animal sampling. Public-genome ascertainment bias may also contribute, as feline isolates are less frequently available and may be preferentially sequenced when clinically significant or antimicrobial resistant. Overall, the data support enrichment of ST147 and other epidemic clones as the principal explanation for higher MDR prevalence in cats, while veterinary exposure, antimicrobial selection, household ecology and sampling bias remain possible contributing factors.

ST147 is a globally disseminated high-risk *K. pneumoniae* clone associated with MDR, ESBLs and carbapenemases, including NDM-, OXA-48-like and KPC-type enzymes [16]. The identification of a compact cat-enriched ST147 sub-lineage containing closely related dog and human-associated genomes is important. Low recombination-filtered SNP distances, a conserved MDR background and shared plasmid replicon architecture suggest recent common ancestry within a host-spanning ST147 population, rather than sporadic recovery of unrelated animal isolates.

The accessory-genome signal further supports strain relatedness across ST147 isolates. Repeated detection of a conserved MDR genotype, including *bla*_NDM-5_, together with recurrent IncFIB(K), IncFII(K) and IncX3 replicon profiles, indicates retention of a clinically important resistance profile across closely related genomes. This does not prove direct transmission between humans and companion animals, nor resolve directionality. However, the presence of human comparator genomes within or immediately adjacent to the same genomic background supports ecological connectivity between host populations. This is consistent with previous comparative phylogenomic studies which show that ESBL-, AmpC- and carbapenemase-producing *K. pneumoniae* from companion animals can belong to major human-associated MDR lineages and share resistance-associated plasmid backgrounds with human isolates [17].

Companion animals are exposed to antimicrobials that can select for MDR Enterobacterales [35, 36], potentially maintaining MDR *K. pneumoniae* once introduced into animal, household or veterinary-hospital environments. However, carbapenems are generally not licensed or routinely used in veterinary medicine; therefore, carbapenemase carriage in companion animals may likely reflect acquisition of resistant strains or mobile elements from human-associated, environmental or healthcare-linked reservoirs rather than direct carbapenem selection [37]. Overall, the cat-enriched ST147 sub-lineage is not evidence that cats are the source of this clone, but it shows that companion animals can carry closely related, clinically consequential MDR lineages overlapping with human-associated genomic diversity.

Collectively, these findings indicate that companion animals should not be viewed as peripheral or epidemiologically separate from the wider *K. pneumoniae* AMR landscape. Instead, dogs and cats can harbour globally disseminated MDR lineages, including high-risk clones with resistance profiles closely related to those observed in human-associated populations. The data do not establish directionality, source attribution or direct transmission, and interpretation is limited by uneven public-genome sampling and incomplete clinical metadata. Nevertheless, the combination of ST overlap, MDR clone enrichment, colonisation-associated recovery and fine-scale ST147 relatedness supports the inclusion of companion animals in targeted One Health genomic surveillance.

## Author Contributions

Stephen Fordham: Conceptualization (lead); writing - original draft (lead); formal analysis (lead); writing - review and editing (lead); Visualization (lead). Elizabeth Sheridan: Conceptualization (supporting); Writing – review and editing (equal). Francis Drobniewski: Conceptualization (supporting); Formal Analysis (supporting); Writing – review and editing (equal).

## Acknowledgments

No funds, grants, or other support were received for conducting this study or preparing this manuscript.

## Conflict of Interests

The authors declare no conflicts of interest.

## Data availability Statement

The data supporting the findings of this study are available in the supplementary material. The human comparator dataset is available at the GitHub repository cited in the Methods.

## Ethics statement

Ethical approval was not required because this study used publicly available bacterial genome sequences and associated metadata and did not involve new sampling of animals or humans.

## Notes

### Competing Interest Statement

The authors have declared no competing interest.

